# Can imputation in a European country be improved by local reference panels? The example of France

**DOI:** 10.1101/2022.02.17.480829

**Authors:** Anthony F. Herzig, Lourdes Velo-Suárez, Frex Consortium, FranceGenRef Consortium, Christian Dina, Richard Redon, Jean-François Deleuze, Emmanuelle Génin

## Abstract

France has a population with extensive internal fine-structure; and while public imputation reference panels contain an abundance of European genomes, there include few French genomes. Intuitively, using a ‘study specific panel’ (SSP) for France would therefore likely be beneficial. To investigate, we imputed 550 French individuals using either the University of Michigan imputation server with the Haplotype Reference Consortium panel, or in-house using an SSP of 850 whole-genome sequenced French individuals.

With approximate geo-localization of both our target and SSP individuals we are able to pinpoint different scenarios where SSP-based imputation would be preferred over server-based imputation or vice-versa. We could also show to a high degree of resolution how the proximity of the reference panel to a target individual determined the accuracy of both haplotype phasing and genotype imputation.

Previous comparisons of different strategies have shown the benefits of combining public reference panels with SSPs. Getting the best out of both resources simultaneously is unfortunately impractical. We put forward a pragmatic solution where server-based and SSP-based imputation outcomes can be combined based on comparing posterior genotype probabilities. Such an approach can give a level of imputation accuracy markedly in excess of what could be achieved with either strategy alone.

## Introduction

Population-based genotype imputation remains a widely used technique for enriching datasets of genotyped or low-coverage sequenced individuals. Advances in software capabilities have been rapid, enormous haplotype reference panels have been assembled, and dedicated computation servers have been created at the University of Michigan^1^ and at the Sanger institute^2^.

Numerous studies have compared the effectiveness of different imputation strategies. The important point of consensus being that imputation benefits from a reference panel that is both large and diverse^2–6^. Public reference panels widely used for imputation include the 1000 Genomes Project (1000G)^7^, the HRC panel^2^ and the TOPMED panel^8^. The size and variety of origin of the reference haplotypes in such panels aims to ensure accurate imputation of target-individuals from different populations. Many groups have published results underlying the importance of preferring ‘local reference panels’ or ‘study specific panels’ (SSPs) – the intuitive concept being that the best panel for imputation should contain reference haplotypes that closely resembles the target individuals. Furthermore as rarer genetic variants are often younger^9,10^, they are expected to be geographically clustered and hence only successfully imputed with geographically relevant reference haplotypes. Increased imputation accuracy coming from SSPs has been shown in populations such as the Netherlands^11^, Estonia^12^, Norway^13^, and Japan^14^. SSP imputation also improves the power of genome-wide association studies (GWAS) involving both common and rare variants^13,15–17^. The benefits of using SSPs have been shown to be particularly evident in the context of isolated populations^17–21^.

SSPs may often be relatively small and so the best approach may often be to combine an SSP with a large cosmopolitan reference panel. Though combining public and study specific reference panels is computationally feasible, it remains problematic for other practical reasons. Panels such as the HRC^2^ or TOPMED^8^ are only fully available through online servers and hence it is not possible to merge their data with one’s in-house sequencing data. Hence, most published results cited above involving a combination of panels have merged an SSP with the freely available (but smaller in comparison) 1000G.

Leading population-based imputation software invoke haplotype copying models based on the Li-Stephens model^22^. This model uses coalescent theory, capturing the idea that if two chromosomes (at a given position) are followed back in time, they will eventually coalesce, sharing a (most recent) common ancestor and this will translate into stretches of shared haplotypes between individuals. For two unrelated individuals, any given genomic region would likely contain many differences representing a very long coalescent time between the pair. But with a large enough sample of a population and in a given genomic region, each observed haplotype can be expected to have a shared lineage (and hence have a relatively recent common ancestor) with at least one other haplotype in the sample. Thus these two haplotypes would likely share a near identical haplotype (allowing for only a few very recent mutations) that would stretch far enough to contain multiple common genetic variants. Extending this idea across regions, a given chromosome from the sample can be described as a mosaic of small haplotype segments present in the pool of all other chromosomes in the sample. This concept is harnessed by imputation software; each target individual chromosome is modelled as a mosaic of reference panel haplotypes using genotyping information for the target individual on a set of common variants. Once a likely chain of copying haplotypes is estimated based on similarities for common genetic variants, missing genotypes can be inferred. Or more often, posterior probabilities of missing genotypes across many potential chains are estimated. Developments in imputation software have been driven by the need to make inference from larger and larger reference panels, but also to operate efficiently to find the best subsets of reference individuals for each chromosomal region. In particular, the PBWT^23^ algorithm has allowed for very rapid sub-selections of reference panel individuals to serve as region-specific reference haplotype pools. PBWT can be employed as a phasing and imputation software on its own but the algorithm has also been incorporated into other software such as EAGLE2^24^, IMPUTE5^5^ and SHAPEIT4^25^. With the concepts of the Li-Stephens model in mind, it is intuitive that imputation will be successful if the reference panel contains relevant haplotypes which closely match the target individual but also enough diversity to enable good haplotype matching across the target’s whole chromosome - i.e. there are no weak links in the chain. This can explain potentially counter-intuitive results such as the inclusion of the UK10K^26^ imputation panel improving the imputation of Italian^27^ and even Chinese^28^ genomes.

Aside from choice of reference panel, an important consideration is the estimation of haplotypes - referred to herein simply as ‘phasing’. The accuracy of phasing has also been widely evaluated, with a parallel rapid development of competing software. Population based phasing software use broadly the same haplotype copying models as imputation software, only that two chains of mosaics have to be found simultaneously rather than a single one. An important difference is that when phasing, inference is often made between individuals in the study. Conversely when imputing, each target individual has missing genotypes imputed from their pre-phased data using only the reference panel. Older software versions such as IMPUTE2^3^ and MaCH^29^ provide the possibility of phasing and imputing simultaneously. Avoiding pre-phasing has been shown to give small increases in imputation accuracy though this comes at a price of a huge increase in computation complexity^30^. Therefore, this approach is unlikely to be considered for imputation involving large target and/or reference panel sample sizes (such as those analysed here); and in particular is not possible on current imputation servers.

Imputation accuracy has not been investigated in French populations. The French population has considerable internal diversity^31,32^ and does not have direct representations in panels such as 1000G, HRC, or TOPMED. Recently, 856 French individuals were whole-genome sequenced at 30-40×, this makes up the FranceGenRef panel (Labex GENMED http://www.genmed.fr/); an obvious candidate for an imputation panel for French genomes. However, as FranceGenRef is relatively small, it is unclear as to whether it will be competitive with a panel such as the HRC (38,821 individuals) for imputation. Furthermore, FranceGenRef does not include individuals from all corners of France and so may not be appropriate for imputing missing genotypes for all French genomes. In this study, we will evaluate potential approaches for both phasing and imputation of French data using either the Sanger and Michigan imputation servers or in-house phasing and imputation. We will also analyse the interplay of population structure within France and the impact that this can have on phasing and imputation accuracy.

## Results

### Evaluating Imputation Servers

Our study involves two French datasets: FrEx, a panel with exome data on 574 individuals recruited in six French cities and FranceGenRef (FGR) with whole genome sequence data on 856 individuals with ancestry in different French regions (Figure 1). The constitutions of both datasets are described fully in the Methods. To motivate the use of a French SSP for imputation of French genomes, an initial investigation of the performance of imputation servers for French individuals was performed. Our technique was to send sets of common variants extracted from FGR to two imputation servers (Michigan and Sanger) in order to be imputed with the HRC reference panel. We could then compare imputed genotypes to the sequence data in FGR. In order to assess imputation accuracy we calculated the imputation quality score (IQS)^33^ per individual, assuming that the true genotypes were those from the sequence data. We established that using the Michigan server and the list of positions on the UK Biobank imputation array provided the most accurate imputation (Supplementary Materials, Supplementary Figure 1). Furthermore, between the different imputation pipelines that we tested, there were always strong correlations between individual IQS statistics (Supplementary Figure 2); the same individuals were always imputed the best (or worst) across our sample. This suggested that underlying characteristic of each individual were determining their individual IQS score (relative to the rest of the sample). A likely cause would be fine-scale population stratification within the sample. To show this, we plotted individual’s IQS scores against their geographical location in France, and a striking pattern emerged (Figure 1); individuals from the North and West of France were imputed with greater accuracy (top deciles of individual IQS scores). The HRC panel contains many individuals of Northern European and Britannic ancestry, this likely explains the higher imputation accuracy for individuals from the North and West of France. This suggests that the internal population structure of France may have a strong influence on the quality of imputation that can be achieved with certain reference panels. Furthermore, using only a panel such as the HRC for imputation in France could lead to an unwanted confounding between imputation accuracy and internal population stratification.

**Figure 1.**
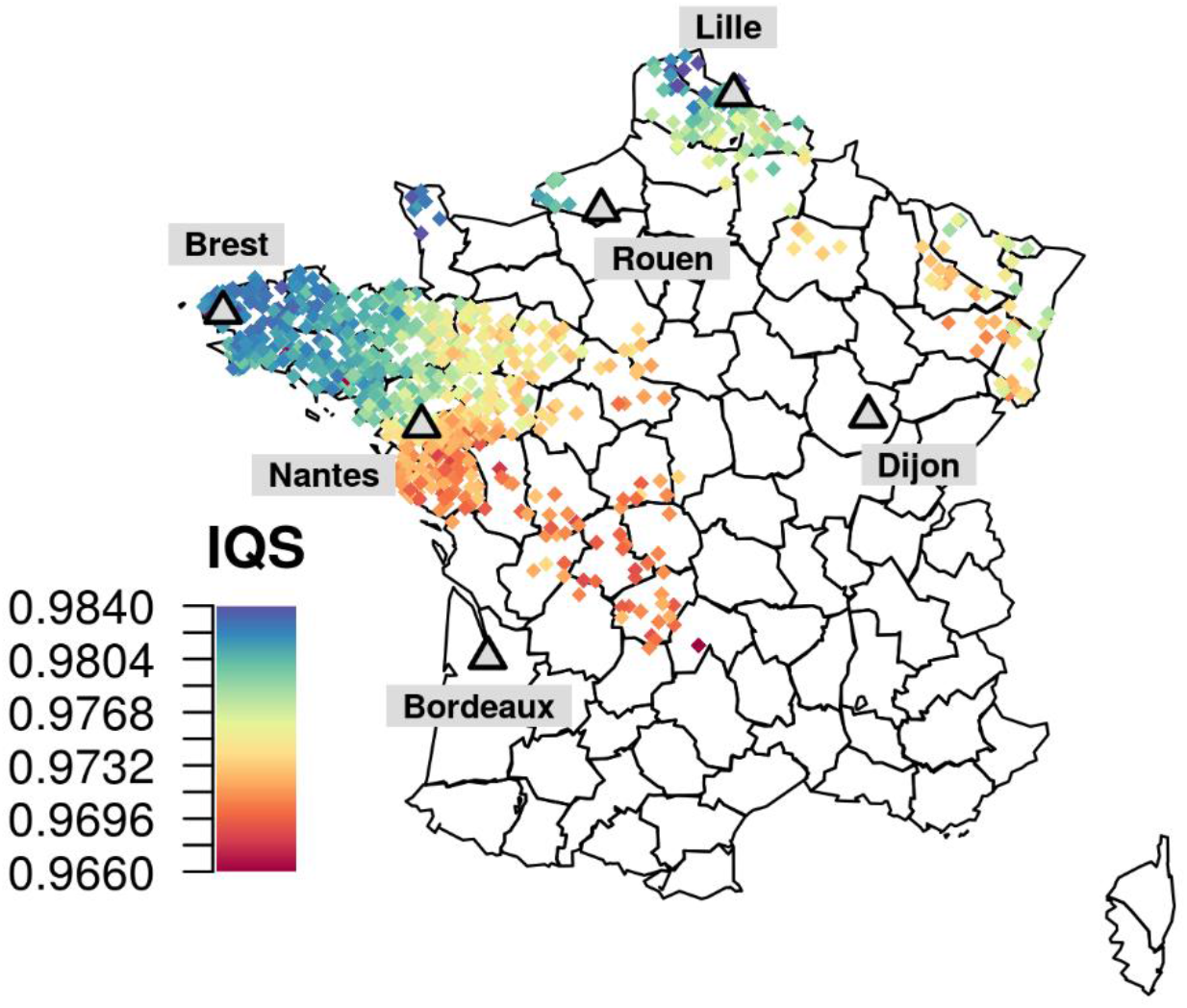
Geographical localities of the participants of FGR (diamonds) and FrEx (grey triangles). For individuals in FGR, positions were estimated as the mean latitude and longitude of the four birth-places of their four grand-parents. For FrEx, recruitment was centred around 6 cities in France (Brest, Nantes, Bordeaux. Rouen, Lille, Dijon). The individuals in FrEx are assumed to have origins close to their recruitment centre as information regarding the origins of each individual’s recent ancestors were used to select participants. Individual IQS scores in FGR from imputation using the Michigan server and the HRC panel are represented by colour. Individual IQS scores range from 0.9661 to 0.9836 (scores closer to 1 represent greater imputation accuracy). IQS was measured on over 17 million variants, a difference in 0.01 between two individuals’ IQS scores approximately represents a swing of 100,000 more or less correctly imputed genotypes.

### Testing different reference panels

To test the impact of using different imputation reference panels in France, we enlisted data from the French Exome Project (FrEX). Here, 550 individuals were analysed (see Methods) who have whole-exome sequencing (WES) data and also genotype data from Illumina OmniExpressExome arrays. We took the array data for FrEx as a basis for imputation and used the WES data for calculating the accuracy of imputation (IQS scores). We sent the array data from FrEx to the Michigan imputation server to be imputed with the HRC panel using the phasing algorithm EAGLE2 and imputation software MINIMAC4. We were also able to effectively use the Michigan server to perform imputation of FrEx using the WGS data of our SSP. This was achieved through the *docker* provided by the Michigan server, allowing us to run their exact phasing and imputation pipeline using our WGS data from FGR as a reference panel whilst being required to send out our WGS panel overseas.

When using the Michigan imputation server, the HRC clearly outperformed FGR (Figure 2, comparing far-left and far-right boxplots MICHIGAN:FGR:FGR against MICHIGAN:HRC:HRC, the notation of each strategy is Place:PRP:IRP where Place (MICHIGAN, SANGER or LOCAL) refers to where the imputation took place, PRP refers to the phasing reference panel, and IRP to the imputation reference panel). Results are split among the six French cities of FrEx. As the HRC panel contains many more individuals than FGR, far more variants can be imputed. The HRC was able to impute 12.6 million variants genome-wide with an RSQ score > 0.5 (the RSQ is the imputation quality score provided by MINIMAC4), compared to 5.2 million variants with an RSQ > 0.5 by FGR. This is due to the fact that our SSP only contains variants with an observed Minor Allele Count (MAC) ≥ 5 in FGR. The superiority of the public panel over the SSP contrasts against many of the results presented in our literature review where SSPs were regularly shown to be the most effective imputation panels. Possible explanations include the small size of FGR and that previous studies had often compared SSP imputation against imputation using the 1000G. But also, this could be partly due to the use of the imputation servers. Examining the described pipeline of the Michigan server, we observed that the phasing step could be working at a disadvantage to the SSP strategy. EAGLE2 is able to very efficiently take advantage of the HRC as a huge phasing reference panel (PRP). However, it has been shown to be less optimal for taking advantage of within-sample phasing^19^, relying more on comparing each haplotype separately to the PRP, which could have had an impact during both the phasing of FrEx and FGR when using the Michigan imputation server.

**Figure 2:**
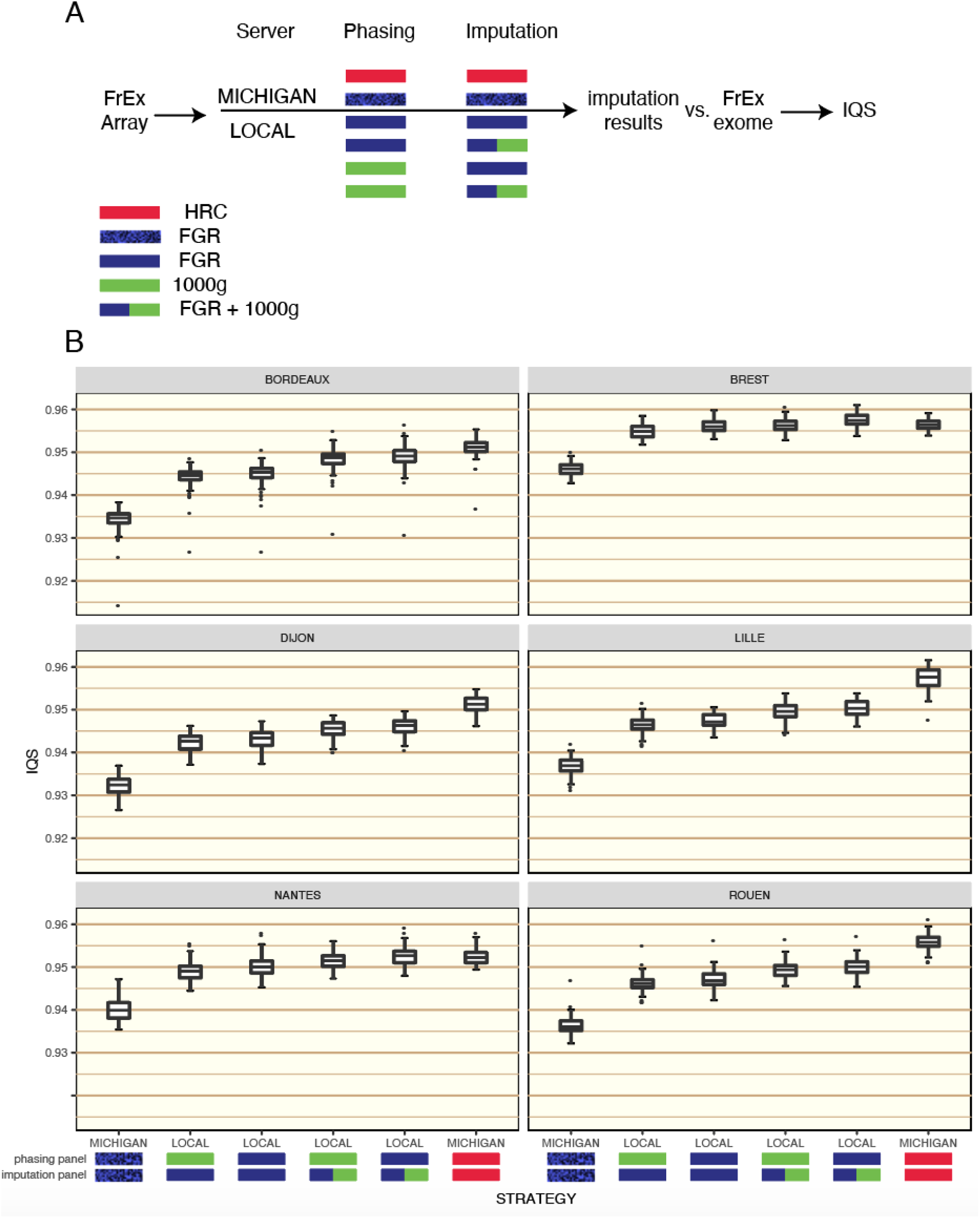
Individual IQS scores for individuals in FrEx for different pipelines. Results are split between the 6 cities of FrEx. Section A) depicts the different possible phasing and imputation strategies that were tested, running either on the Michigan server or locally (LOCAL) in our lab and with different combinations of phasing and imputation panels. Section B) gives boxplots of individual level IQS scores for each strategy. Of the 550 individuals analysed, 89 are from Bordeaux, 96 from Brest, 87 from Dijon, 93 from Lille, 90 from Nantes, and 95 from Rouen.

We constructed our own phasing-imputation pipeline to assess the impact of using the imputation servers (description in Methods). To further investigate the impact of the pre-phasing step, we phased FrEx using SHAPEIT4 with either FGR or 1000G as a PRP. By using IMPUTE2 with the merge-ref-panel option, we were able to use a combined reference panel of the 1000G and FGR without having to restrict to a common list of variants. When the PRP and IRP were FGR, the imputation quality was improved substantially by using our in-house imputation pipeline compared to the results achieved with the Michigan server via the docker (LOCAL:FGR:FGR against MICHIGAN:FGR:FGR). In the cities of Brest and Nantes, SSP-based imputation became competitive with the server-based approach using the HRC panel. These two cities lie in the regions that are most well represented by FGR. For all six cities, a marginal improvement in imputation accuracy was gained by including the 1000G in the reference panel (LOCAL:FGR:FGR against LOCAL:FGR:FGR+1000G).

Regarding the key comparison of imputation results between LOCAL:FGR:FGR and MICHIGAN:HRC:HRC, we found that the advantage brought by the HRC panel over FGR was particularly evident for rare variants (see Supplementary Figure 3 where the mean IQS scores per-variant were compared in detail between these two strategies for different categories of MAF). Only in Brest and Nantes did the two pipelines perform similarly. However, this analysis does not tell the full story as IQS was calculated on only variants that can both be imputed by the HRC and our FGR. Indeed, this ignores the existence of potentially-population specific variants that cannot be imputed by panels such as 1000G or the HRC. In Supplementary Figure 4, we show the proportions of variants observed in FrEx that are also observed in the other three datasets (FGR, 1000G, HRC) pertinent to this study. For the rarer variations in FrEx, large proportions of variants are not observed in all three of FGR, 1000G, and HRC including many that are only observed in FrEx and FGR. This highlights the importance of including an SSP (ideally in conjunction with a public panel) in order to impute such population specific variants.

### Investigating haplotype sharing between target and reference individuals

As Brest and Nantes are located in the regions that are the best represented in France by FGR, it is likely that the FrEx individuals from those two cities exhibit greater haplotype-sharing with FGR than FrEx individuals from the other four cities. Hence, we investigated the composition of the estimated haplotypes in FrEx to reveal where our imputation pipeline was most accurate. As imputation is haplotype based, the haplotype-estimation pre-phasing step clearly has an impact on the quality of imputation. We postulated that using an SSP as PRP could be particularly beneficial as haplotype estimation could be improved and imputation using the same SSP would thus be facilitated. This we demonstrated by comparing phasing-imputation run LOCAL:FGR:FGR against LOCAL:1000G:FGR. To approximate the accuracy of phasing, we applied the principal of phasing uncertainty. This involved repeatedly performing phasing using different random seeds and evaluating the stability of the final estimated haplotypes. Using such a method, we could have an approximation of the Switch Error Rate (SER) of the haplotypes constructed (see Methods). We refer to the two different phasing outputs involved in these two pipelines as FGR-PRP and 1000G-PRP.

Individual phasing uncertainty statistics were strongly correlated with the eventual IQS scores (Supplementary Figure 5 - left panel) which demonstrates that the phasing uncertainty statistics calculated have likely captured a true approximation of the phasing quality. It was observed that the level of improvement in SER and IQS coming from using FGR as a phasing reference panel was noticeably higher for the two cities of Brest and Nantes compared to the other four cities of FrEx (Supplementary Figure 5 - right panel).

Furthermore, we explicitly looked for regions of haplotype-sharing between FrEx individuals and FGR individuals by estimating IBD segments using RefinedIBD^34^. We clustered individuals in FGR based into 12 groups (see Methods) using finestructure^35^. These coincided with different geographical regions of France (Figure 3). We calculated the total IBD shared between each city in FrEx and each cluster of reference haplotypes in the two different phasing scenarios described above: FGR-PRP and 1000G-PRP. Far greater total shared IBD was estimated between Individual’s from Brest and Nantes and FGR under FGR-PRP. What is more, the increase in detectable haplotype-sharing pertained largely to shared segments of length greater than 3cM between individuals of Brest and Nantes in FrEx and individuals in FGR from the regions of France close to Brest and Nantes. For the detection of shorter segments, the choice of PRP had less impact.

**Figure 3.**
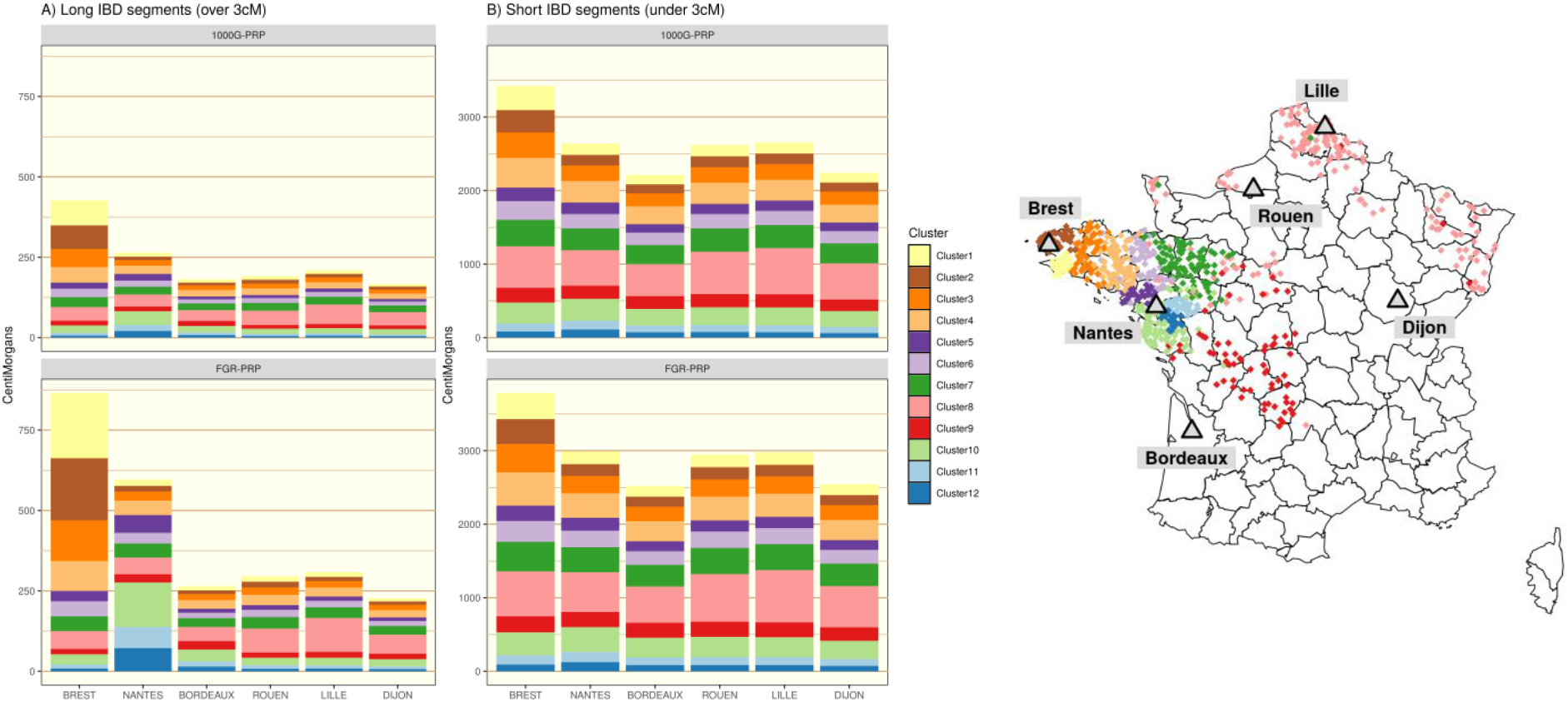
Left: Haplotype sharing between individuals from the different FrEx cities and individuals from FGR. Results are split by the 12 clusters of FGR detected with finestructure. The PRP used in the phasing step was either with 1000G (top panel 1000G-PRP) or FGR (bottom panel FGR-PRP) and IBD segments were split into long segments over 3 cM (A - left column) or small segments under 3 cM (B - right column). Right: A map of France showing the 12 haplotype-sharing clusters in FGR with the 6 cities of FrEx highlighted (grey triangles). Colours of each haplotype-sharing cluster correspond to those in the plot on the Left.

To demonstrate the interplay of the estimation of shared IBD segments between the target and reference panel and imputation accuracy, we examined the imputation of rare variants (determined by minor allele frequency less than 0.01 in FrEx) for the FrEx individuals from Brest. Imputation accuracy was evaluated in two variant sets: those inside long IBD segments and those outside long IBD segments. Specifically, for each rare-variant, we tabulated the imputed dosages of heterozygous sites observed in the individuals of Brest inside and outside long IBD segments (>3cM) shared with Clusters 1 and 2 of FGR that cover the Finistere department where Brest is located. Histograms of these imputed dosages (which should be equal to 1 if the imputation has been successful at a heterozygous site) are presented in Supplementary Figure 6 for two imputation strategies: LOCAL:FGR:FGR and MICHIGAN:HRC:HRC. There was a clear higher proportion of correctly imputed genotypes for variants within IBD segments compared to variants outside IBD segments under the LOCAL:FGR:FGR imputation strategy but not under the MICHIGANC:HRC:HRC strategy. This shows how imputation using a local reference panel can be expected to improve accuracy, and in particular for the rare local variants that are expected to lie on long IBD segments. This is because rare variants are expected to be younger than common variants so if it is given that two individuals share a haplotype containing a rare allele, the haplotype is expected to also be relatively young and thus relatively long.

### Pragmatic imputation strategy for France

We have thus far demonstrated that by optimising certain stages of our phasing and imputation strategy we could achieve a comparable imputation quality using FGR compared to imputation servers that have access to the HRC panel. However, this was only the case for the individuals of Brest and Nantes in FrEx. Furthermore, it was also observed through analysing haplotype-sharing that even for Brest and Nantes the SSP imputation was not performing uniformly and that its accuracy would vary in different genetic regions. Ideally, a combined panel of FGR and HRC could be used. Without a straightforward path to this solution, one possibility is to attempt to combine imputed data from two different imputation runs in the spirit of the PedPop method put forward by Saad & Wijsman^36^. This simply involves merging two or more imputation outputs such that all variants imputed by any of the strategies are present. For variants that are imputed by multiple imputation strategies, a simple ‘most confident vote’ selection is used (see Methods). We combined imputation using the MICHIGAN:HRC:HRC with our own LOCAL:FGR:FGR imputation in such a manner (see Methods) and the overall improvement to the imputation accuracy was substantial (Figure 4). This hybrid imputation coming from this combination is denoted as HYB.

**Figure 4 -.**
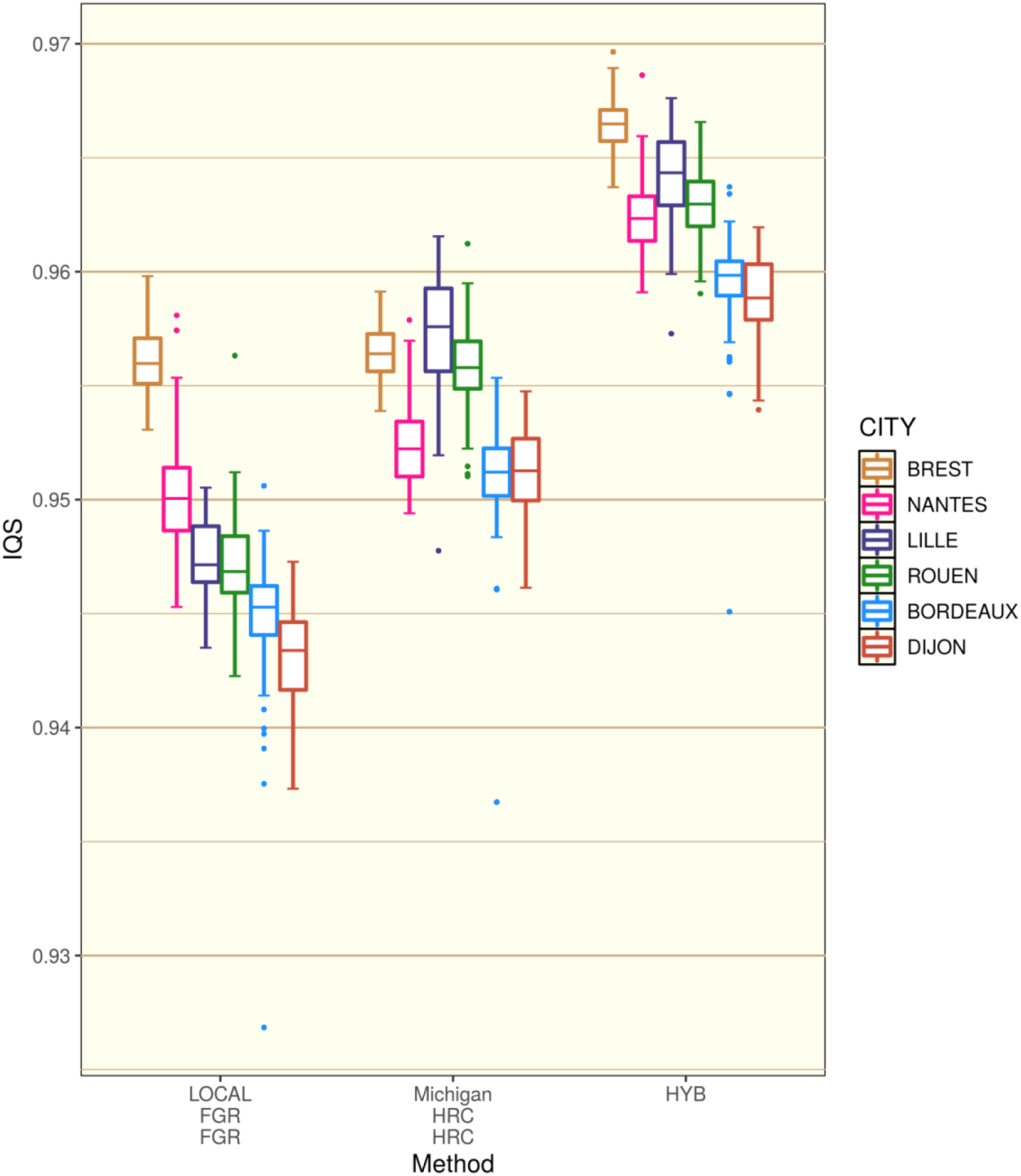
Individual IQS scores for the hybrid (HYB) imputation strategy split by city in FrEx against the previously calculated scores for strategies LOCAL:FGR:FGR and MICHIGAN:HRC:HRC described in Figure 2.

To illustrate how the HYB method was improving the imputation, for each individual, IQS scores were calculated for two sets of variants: One set where there was agreement between the two imputation strategies regarding the most likely genotype (Accord), and a second where there was disagreement (Discord). The average percentages of genotypes in agreement for each individual for the six cities of FrEx were: Bordeaux 98.5%, Brest 98.7%, Dijon 98.5%, Lille 98.6%, Nantes 98.6%, and Rouen 98.6%. Overall, agreement between HRC and FGR imputation corresponded with the correct genotype being assigned the highest probability 98.5% of the time. Hence, agreement between HRC and FGR was a reliable indication of accurate imputation. Choosing the set of imputation probabilities with the greatest top probability in the case where the two imputation runs are in Accord will therefore produce a dosage closer to the true genotype for the majority of cases. This gave a significant boost to the IQS statistics (Figure 4, Supplementary Figure 7) and would also lead to an increase in power for prospective association tests. In Supplementary Figure 7, we also observe that this improvement afforded by the HYB strategy was present for both rare and common genetic variants.

The two imputation runs were not in agreement (Discord) for 1.5% of all genotypes analysed (125,442 exonic variants). This corresponds to approximately 1800 genotypes per individual. In this set, the percentage of variants where HRC imputation was correct was the following for the different cities: Bordeaux 64%, Brest 59%, Dijon 66%, Lille 69%, Nantes 61%, and Rouen 67%. The hybrid imputation strategy chose the correct genotype more often than not (Bordeaux 68%, Brest 68%, Dijon 67%, Lille 70%, Nantes 68%, and Rouen 69%). Hence, the HYB strategy coped well with disagreement between the two pipelines. In Brest and Nantes, HYB even provided an improvement in imputation in the Discord set of variants (Supplementary Figure 7); both for rare and common variants. Therefore, a simple combination of in-house and server-based imputation could provide a pragmatic and most effective imputation strategy.

## Discussion

Many factors affect the accuracy of phasing and imputation, though most focus has been put on the size and composition of the haplotype reference panel. Furthermore, imputation has shifted from an operation performed in-house using publically available data and free academic software to an operation that is increasingly performed at distance using publically accessible imputation servers. The main motivations for using external imputation servers is their convenience and the access they provide to the largest (and hence most powerful for imputation) public reference panels. However, certain compromises currently have to be made if one decides to use an imputation server. Importantly, there is not the possibility to combine an SSP with public reference panels, and there is less control of haplotype phasing. In this study we have shown that such considerations can make a significant difference for imputation quality. Hence, the optimisation of an imputation strategy goes far beyond simply choosing the largest available reference panel. If an SSP is to be used, we suggest that at this current time it is still preferable to perform phasing and imputation in-house rather than turning to online imputation servers.

We decided that, for this study, it was not necessary to test the very recently developed TOPMED server. This was for two reasons, firstly the TOPMED panel is aligned to genomic build 38 and it was beyond the scope of this study to re-call the FranceGenRef and FrEx datasets that are both aligned to build 37. Secondly, whilst the TOPMED reference panel is significantly larger than the HRC, it contains a comparable number of individuals of European ancestry. The TOPMED panel has been shown to greatly outperform the HRC for imputation of individuals of Latin American and African ancestry^8^ but would not provide such a significant improvement in accuracy compared to the HRC for French individuals.

It would have been possible to gain permission from the European Phenome-Genome Archive to download a part of the HRC panel (https://ega-archive.org/studies/EGAS00001001710). This option was used in one simulation study^19^, where this subset of the HRC was combined with an SSP for evaluation of imputation in an isolated population. Using this subset of the HRC has recently been shown to be effective for imputation in conjunction with SSPs by Quick et al.^37^ We decided not to pursue this avenue in this study for two reasons. Firstly, having to download this subset of the HRC and then perform imputation using IMPUTE2 and the merge-ref-panel option is very computationally heavy, requires a lot of storage space, and requires the submission of a specific request for the HRC subset. Hence, this is a strategy that may not be suitable for all researchers and so is not a realistic recommendation to make. Secondly, in this study we wished to focus on the pros and cons of the choice of using the relatively easy server-based approach against in-house imputation. Hence, to test the HRC we preferred to access it through the server; furthermore, this allowed us to test the HRC panel in its entirety.

Our results chime with previous results regarding the benefits of local reference haplotypes^11,12,14–16,18,20^. However, by including the complete HRC panel in our study, we showed certain limitations to SSP-based imputation. This was possible by investigating the fine genetic structure in France and its impact on the imputation of French genomes. Our SSP was successful in improving imputation beyond the possibilities of the HRC panel but only for target individuals that were from the regions that were densely covered in FGR (Brest and Nantes). In the other four cities, the HRC clearly afforded higher accuracy. We note that evaluating imputation accuracy per-individual rather than the more commonly used per-variant calculation was important in uncovering such patterns. Indeed, imputation in France using only the HRC led to a clear gradient of imputation quality in FGR. This further motivates the use of local reference haplotypes to avoid the potential of introduced bias as panels such as the HRC will likely provide stronger imputation for individuals from the North and West of France.

We have also demonstrated that the benefits to SSP-based imputation coincided with the sharing of haplotypes between the target and reference individuals. This highlights the importance of optimising haplotype estimation, an area we have concentrated on in this study. Indeed, the imputation of rare-variants would likely benefit noticeably from greater accuracy in the phasing of the SSP. Further improvements to phasing performance could also be sought either through read-based phasing algorithms^38^ or through consensus based phasing^39,40^. Another promising approach is to replace array based genotyping with low-pass sequencing^41,42^.

The FGR panel used here contained 850 individuals. The largest prospective SSP for France, the POPGEN project of the French medical genomics initiative^43,44^ will contain roughly 4,000 individuals. Joining the dots, the significant improvements to the estimated SER and IQS for the individuals of Brest and Nantes would suggest that imputation could be highly accurate for individuals from across France using this novel reference panel. Particularly as we observed that the HRC panel performed less well for individuals from towards the South of France, an area that will be well represented in POPGEN. However, there may still be room to incorporate imputed variants from imputation servers due to the undeniable power of huge public reference panels for imputation; in particular, for rare variants. Rare variants that arrived recently in the population can be expected to have a high level of IBD-sharing^45^ lying within long shared segments. Such variants should be expected to be well imputed using an SSP. This does not hold for older variants that are observed to be rare due to many generations of purifying natural selection^46^; for such variants, the breadth provided by large cosmopolitan panels may provide the best imputation. The POPGEN dataset will cover the whole of France but realistically may not provide a significantly denser coverage than what is given by FGR for the regions of Bretagne and Pays-de-la-Loire (the regions that surround Brest and Nantes, respectively). Combining panels allows for a greater number of overall variants to be imputed as panels will not have coinciding lists of observed variants; each will have a set of variants only observed in that panel. Without the current possibility to combine an SSP with the full HRC or TOPMED, we have put forward a simple pragmatic approach for combining in-house and server-based imputation in order to give the more complete and accurate imputation.

## Methods

The two sequencing datasets used in this study, FrEx and FGR were prepared using VCFprocessor^47^ using in-house settings (described in Supplementary Material). Individual were selected for FranceGenRef using strict criteria on ancestral places of birth. Specifically, individual were only sequenced if their four grand-parents were known to have all been born near to one another. The exact criterion was that all pairs of grand-parents would be born within no more than 30 km of each other. By taking the barycentre of the co-ordinates of all 4 grand-parents, we approximated the ancestral location of each individual in FGR. 862 individuals were recruited from three sources, 458 from the GAZEL cohort (www.gazel.inserm.fr/en)^48,49^, 354 from the PREGO cohort (www.vacarme-project.org), and 50 blood donors from the Finistere region. All individuals signed informed consent for genetic studies at the time they were enrolled and had their blood collected. The recruitment is described in full elsewhere [reference to another FranceGenRef paper submitted at the same time?]. The FrEx data analysed here comprises 557 (out of 574) individuals who are those who have both genotyping (Illumina OmniExpressExome arrays) and WES data. A total of 824,279 variants are found in the WES data after QC. Our SSP was built with 856 individuals with WGS data from FGR. Individuals were excluded from both FrEx (7 removed) and FGR (6 removed) for our study due to either the individuals being present in both panels, to close relatedness with other individuals, or due to quality control measures (see supplementary materials). We kept only variants with a minor allele count above 5 for the creation of an imputation panel. For both datasets, we have approximated geographical locations for each individual. Individuals included in the FREX project are 574 healthy individuals sampled in 6 different regions of France around 6 cities (Bordeaux, Nantes, Brest, Rouen, Lille and Dijon). These individuals are either blood donors with grand-parents born in Finistere or Pays de la Loire (Brest and Nantes samples), or unaffected spouses of individuals affected by Alzheimer’s disease (Rouen and Lille samples) or individuals from two different cohorts (3 cities Dijon^50^ for the Dijon samples and PAQUID^51^ for the Bordeaux samples).

Imputation quality was measured using IQS^33^ calculated per-individual across various sets of genetic variants. This imputation score measures the concordance between the truth set and the posterior imputation probabilities whilst taking into account the expected level of concordance by chance. When splitting results by minor allele frequencies (MAFs), we used the naive MAF estimates from FrEx and results are shown either for rare variants (MAF < 0.01) or non-rare variants (MAF ≥ 0.01). IQS was calculated on a set of 125,442 exonic variants across the 22 autosomal chromosomes that could be imputed with the constructed imputation panel of FGR (i.e. variants that passed the quality control measures in FGR and thus had a minor allele count superior to 5). To describe an imputation strategy, we use the following notation: Place:PRP:IRP where Place refers to the location of the imputation (either Michigan imputation server, the Sanger server, or in-house at LOCAL), PRP refers to the phasing reference panel, and IRP refers to the imputation reference panel.

In order to use FGR as a reference panel for our-in house LOCAL pipeline, it was phased using SHAPEIT4^25^ and the ‘sequencing’ option to optimise the algorithm for WGS data. Furthermore, in an effort to improve the phasing performance of SHAPEIT4, we specified the following iteration programme: ‘8b,1p,1b,1p,1b,1p,1b,1p,15m’. Conversely, when using the Michigan imputation server with the FGR panel for the Michigan:FGR:FGR strategy, the phasing of FGR was performed using the Michigan server and the phasing-only functionality.

FrEx was phased with SHAPEIT4 and imputed using IMPUTE2. The choice of IMPUTE2 may seem questionable given the availability of more recent version such as IMPUTE5^5^ as well as competing software such as MINIMAC4 or BEAGLE5^52^. IMPUTE2 was chosen purely due to the availability of the merge-ref-panel option, allowing for a combined panel of the 1000G and FGR to be used. The importance of this option is demonstrated by the observation that 0.54% of all variants in the SSP we constructed from FGR are not present in the 1000 Genomes Projects. Without this cross-imputation option, these SSP-specific variants would be lost. Given that software cited above rely on similar methodology and have similar performances^5^ (with more recent versions admittedly bringing incremental improvements), we felt that this was a suitably choice for putting forward an imputation strategy involving a SSP. The improvements that have been made to imputation software beyond IMPUTE2 are concerned with the ability to leverage vast reference panels such as the HRC or TOPMED. Our imputation strategy LOCAL:FGR:FGR+1000G involves a combined reference panel of only 6708 haplotypes and so it is reasonable to employ IMPUTE2 in this scenario. However, using IMPUTE2 with a combination of the HRC and our SSP would encounter excessive runtime.

To approximate Switch Error Rate (SER) without knowing the true phase in FrEx, we simply ran SHAPEIT4 21 times using 21 different random seeds. Across the 21 repetitions, and for each pair of adjacent heterozygous genotypes, we assumed that the phase configuration assigned by the majority of random seeds was the correct phase; this allowed us to estimate SER in each seed before finally calculating an average SER across all 21 replicates.

IBD segments in FGR were estimated using RefinedIBD^34^. The resultant matrix of IBD sharing between individuals was then treated as matrix of ‘chunk lengths’ and supplied to finestructure^35^ to establish 12 groups of individuals likely having similar genetic backgrounds. As described in Bycroft et al.^53^, using a chunk-length matrix necessitated the estimation of the ‘c-factor’ parameter from within the sample, for which we followed the instructions given in the supplementary material of Bycroft et al.^53^. The choice of 12 groups was made by inspection and in order to give a set of easily interpretable groups. Up until 12 groups, each cluster identified corresponded to over 10 individuals and to specific geographical region. Beyond 12, groups become small and lacked easily interpretable links to geographical regions. We note that finestructure was unable to distinguish the individuals in FranceGenRef from the North and the East of France. We attribute this to the fact that we don’t have a sufficient sample size in these regions and that, as observed by the Eigen decomposition of the IBD sharing matrix (see Supplementary Figure 8), the most evident sources of variation in the data come from the proximity of individuals to the source populations of the Brittany region and the Pays-de-la-Loire region. As FranceGenRef does not represent a fair sampling of the French population, it is not surprising that the finestructure analysis largely reflects only the variation in the West of France; where we have by far the most individuals. However, the clusters presented here are still relevant for the West of France and are instructive in showing the potential for extensive fine-structure in the French population.

To combine the imputation pipelines for the HYB imputation. We simply compared the maximal probabilities for each pair of genotype from the pipelines MICHIGAN:HRC:HRC and LOCAL:FGR:FGR. For example, if the posterior imputation probabilities for a genotype of a given individual are *I_A_* = (0.95,0.05,0.00) & *I_B_* = (0.85,0.15,0.00) from imputation strategies A and B, respectively, then only the posterior probabilities *I_A_* will be retained as they are the most certain. The concept that the more certain a set of genotype probabilities the more accurate the imputation is well known and underpins the calculation of most imputation quality metrics^54^. Inspection suggested that when the two maximal probabilities were very close, little could be gained by selecting the trio with the highest probability. Furthermore, due to the differences in imputation software (MINIMAC4 against IMPUTE2), we often saw that the maximal probability of LOCAL:FGR:FGR (denoted as 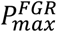) was larger than the its counterpart 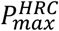 but only by an order of 10^−2^. We found that an effective combination method was to select the imputation trio of posterior probabilities from LOCAL:FGR:FGR if and only if 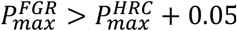, hence minorly giving priority to HRC when the 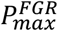 and 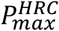 were very close. This rule was used to form the HYB imputation presented in the Results section. Variants were split into groups denoted as Accord and Discord, based on whether 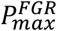 and 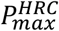 indicated the same genotype or not.

## Acknowledgments

We would like to thank all of the members of the FranceGenRef cohort and the FrEx cohort for their participation in this study. Financial support was obtained from the French Ministry of Research and Innovation for the POPGEN project from the French Medical Genomics Plan. Administrative and regulatory support was provided by Inserm. Funding information: Laboratory of Excellence GENMED, Grant/Award Number: ANR-10-LABX-0013; Funding from the French Ministry of Higher Education, Research and Innovation for the POPGEN France Genomic Medicine pilot project Funding from Inserm Cross-Cutting Project GOLD

## Data Availability

Those wishing to access the data on a collaborative basis should contact Emmanuelle Génin. Summary information for the FrEx dataset is available at http://lysine.univ-brest.fr/FrExAC/.

## Author Contributions

AFH wrote the manuscript and carried out all major analyses. LVS assisted in the analyses, notably in relation to the use of the external imputation servers. All analyses in the study were discussed and decided upon by AFH, LVS, and EG. CD, RR, JFD, and EG contributed to data production for both the FrEx and FranceGenRef datasets. All authors participated in the final redaction of the manuscript.

## Declarations

The authors have no conflicts of interest to declare.

