## Supplementary Materials for "Can imputation in a European country be improved by local reference panels? The example of France"

**1 :** Inserm, Univ Brest, EFS, UMR 1078, GGB, Brest, France
*Faculté de Médecine - IBRBS 22 avenue Camille Desmoulins F-29238 BREST Cedex 3 - France*
**2 :** CHRU Brest, Brest, France
**3 :** LABEX GENMED, Centre National de Recherche en Génomique Humaine, Evry, Paris
**4 :** unité de recherche de línstitut du thorax UMR1087 UMR6291
*Université de Nantes, Institut National de la Santé et de la Recherche Médicale : U1087, Centre National de la Recherche Scientifique : UMR6291*
*8 quai Moncousu - BP 70721 - 44007 Nantes Cedex 1 - France*
**5 :** Centre National de Recherche en Génomique Humaine
*CEA-DRF-IBFJ-CNRGH, GENMED, Fondation Jean Dausset CEPH*
*Institut de Biologie François Jacob, CEA, Université Paris-Saclay, F-91057, Evry - France*

***Details on the utilisation of imputation servers:***

We carried out a set of tests of the performance of two imputation servers (Michigan and Sanger). We were able to vary the choice of variants to represent different possible genotyping arrays (Supplementary Figure 1). We tested the following arrays: The Illumina Core Exome array (ICE), the Axiom Precision Research Array (PRMA), and the UK Biobank imputation array (UKBB). Furthermore, we could also test different imputation software (MINIMAC4[1] and PBWT[2]). For both servers, EAGLE2[3] was used as phasing software as this is currently the only choice available on the Michigan server. These results are presented in Figure 1. We found that the genotyping array that clearly gave the most accurate imputed genotypes was UK Biobank Axiom array. Imputation accuracy was overall slightly higher on the Michigan server compared to the Sanger server, probably due to the difference in imputation algorithm being used (MINIMAC4 or PBWT, respectively).

***Details on Quality Control using VCFProcessor[4] using the QC1078 setting:***

Genotypes were set to missing when:

- Depth (DP) < 10
- Genotype Quality (GQ) < 20

Subsequently, variants were excluded (for all individuals) using the following criteria for various summary statistics (measured across all samples) generated by GATK v.3.8[5].

- Allele Balance for Heterozygous calls (ABHet) outside of the range [0.25,0.75]
- Quality-By-Depth < 2
- MQRankSum < - 12.5 (Z-score from Wilcoxon rank sum test of Alt vs. Ref read mapping qualities)
- Mapping Quality (MQ) < 40 for SNPs or < 10 for INDELS.
- Strand Bias odds ratio > 3 for SNPs or > 10 for INDELS
- Fisher's exact test for strand bias (phred scaled p-value) > 60 for SNPs or > 200 for INDELS
- HQRatio < 0.8
- Inbreeding Coefficient (estimated) < -0.8
- Callrate < 0.9

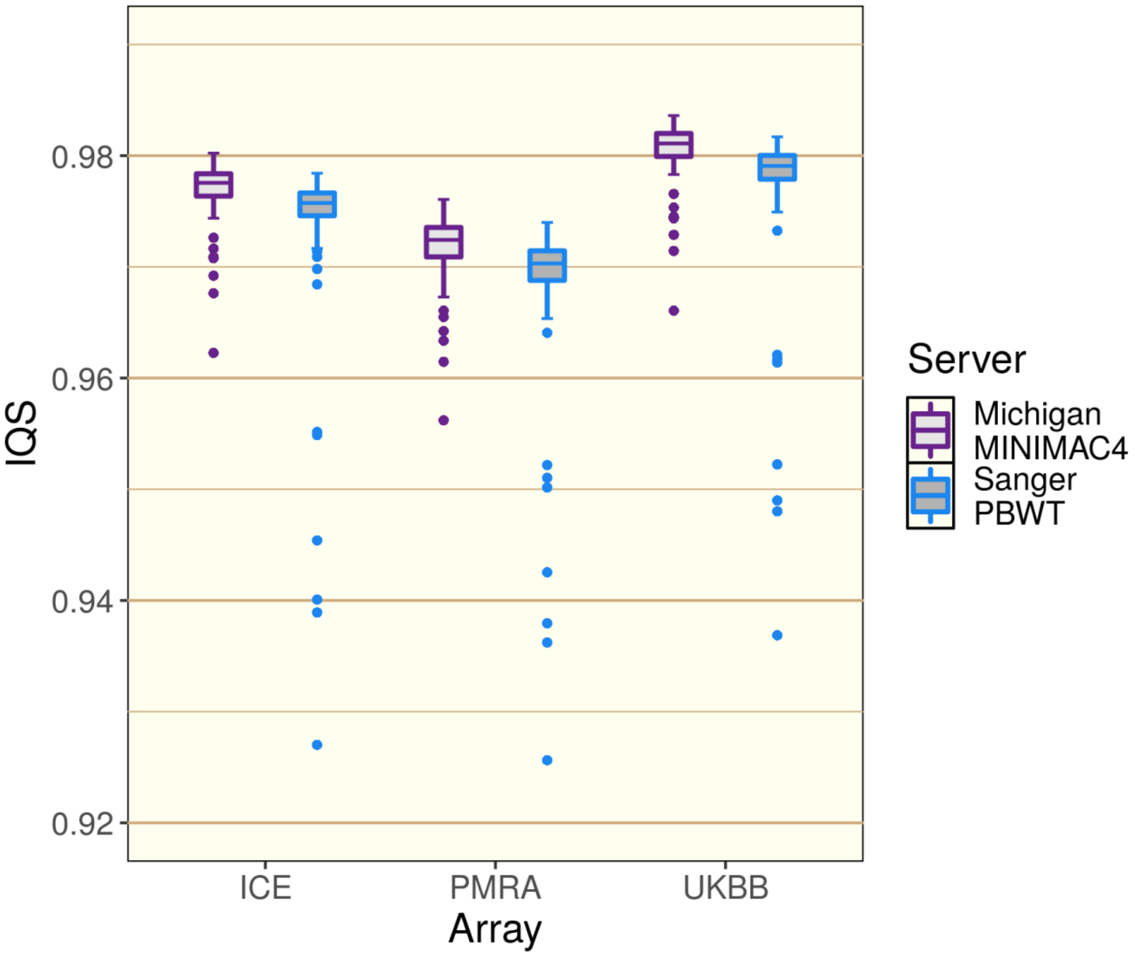


*Supplementary Figure 1: Imputation accuracy of different servers and arrays. For each individual, array positions were extracted, sent to an imputation server before calculating an individual Imputation Quality Score (IQS) by comparing the sequenced non-array positions against imputed counterparts. ICE: Illumina Core Exome array, PRMA: Axiom Precision Research Array, UKBB: UK Biobank imputation array.*

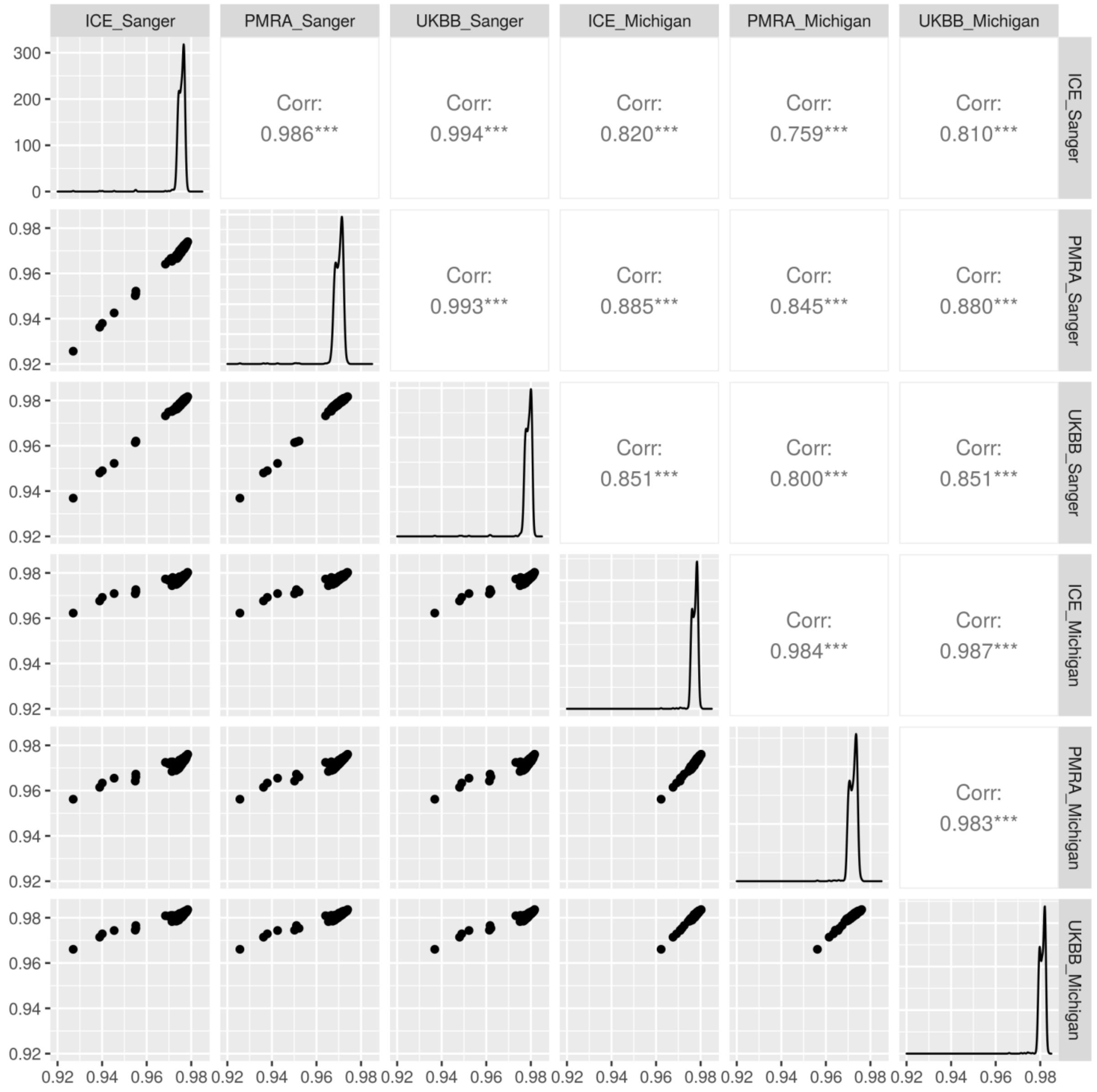


***Supplementary Figure 2: The same individuals are imputed well or poorly, irrespective of the imputation server or array chosen. Corr = Correlation. ICE: Illumina Core Exome array, PRMA: Axiom Precision Research Array, UKBB: UK Biobank imputation array.***

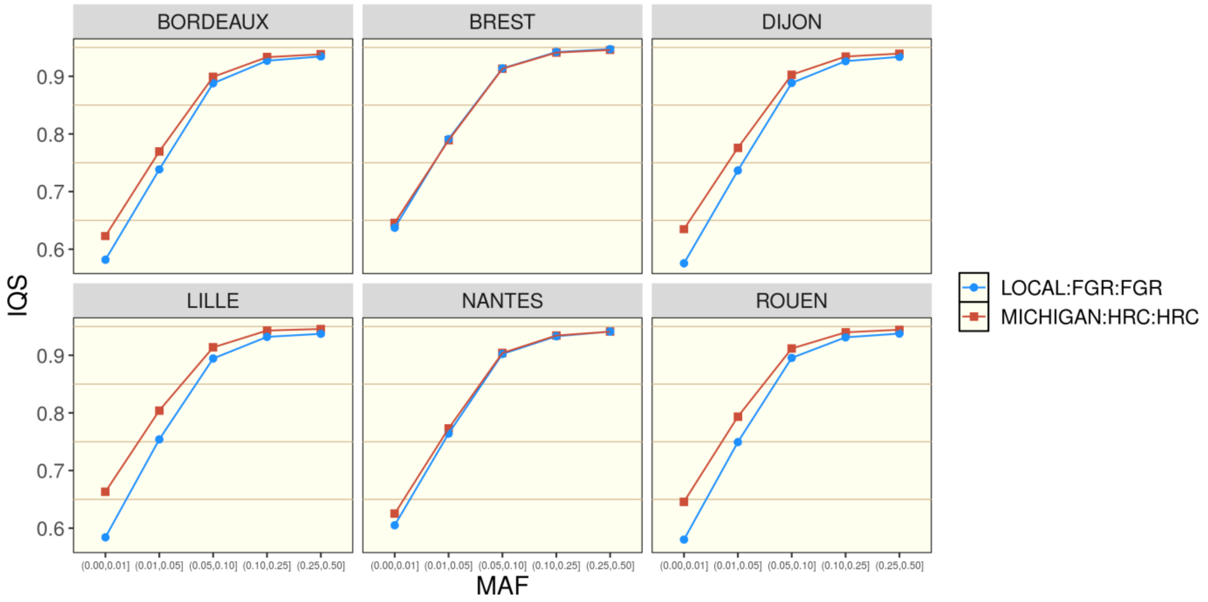


**Supplementary Figure 3 - MICHIGAN:HRC:HRC vs LOCAL:FGR:FGR. Mean per-variant IQS scores for five different minor allele frequencies (MAF) bins are presented. MAF was measured separately in each city by counting the observed number of minor alleles in each group of FrEx.**

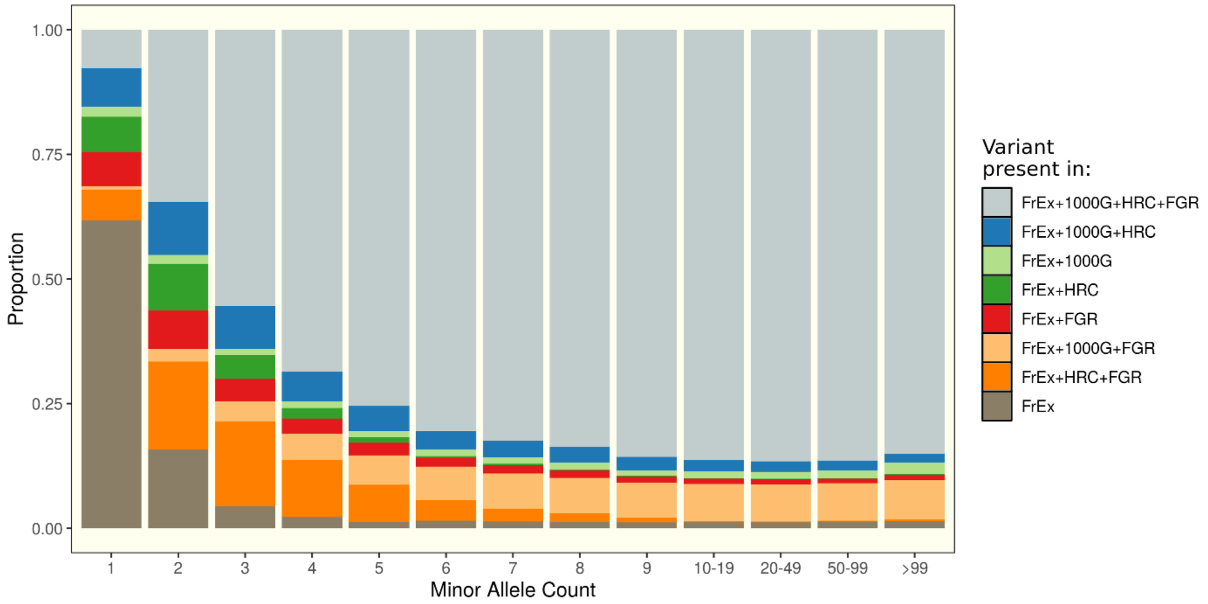


**Supplementary Figure 4. Proportions of variants which are observed in FrEx and other dataset split by different Minor Allele Count (MAC) bins measured in FrEx. For example, the group 'FrEx' represents the variants that are observed in FrEx but nowhere else, 'FrEx+1000G+HRC+FGR' represents the variants observed in all 4 datasets, and 'FrEx+FGR' represents the variants observed only in the French datasets FrEx and FGR.**

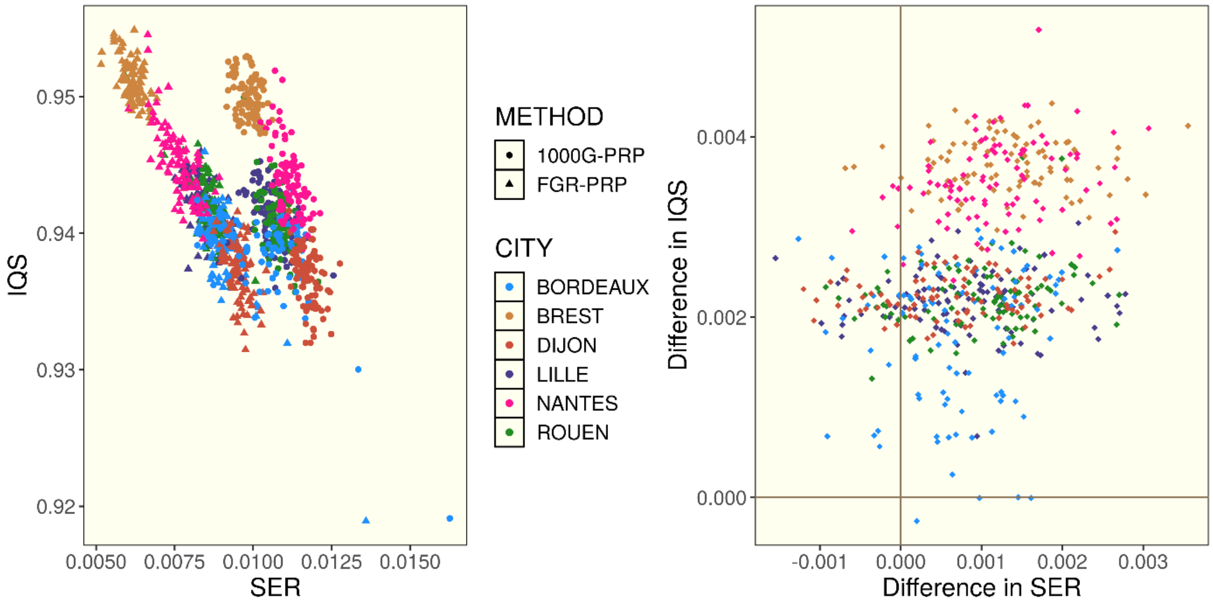


**Supplementary Figure 5.** ***Comparison of imputation and phasing accuracy results using either FGR (FGR-PRP) or 1000G (1000G-PRP) as a phasing reference panel for FrEx. Right: Individuals approximated Switch-Error Rate (SER) statistics against individual Imputation Quality Score (IQS) statistics. Clear correlation was observed in both clouds of points (FGR-PRP or 1000G-PRP). Left; to show that the change of PRP has a more pronounced consequence on the individuals coming from Brest and Nantes, the differences in the SER and IQS statistics from the Right panel are plotted. The individuals from Brest and Nantes have noticeably greater differences (representing greater improvement) in both statistics when moving from 1000G-PRP to FGR-PRP.***

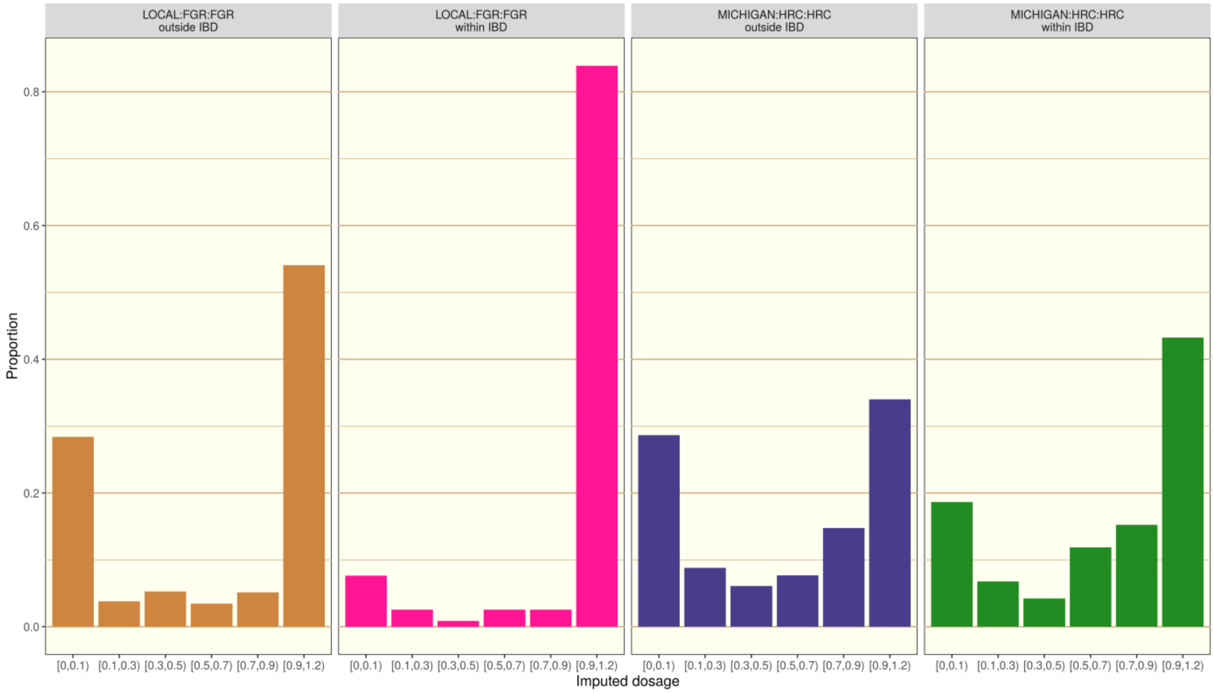


**Supplementary Figure 6 - Imputed dosage of heterozygote genotypes of rare-variants (MAF<0.01) obtained for FrEx individuals from Brest for two different sets of variants, those inside and outside of the long IBD segments shared with FGR clusters 1 and 2 and for two different imputation strategies (LOCAL:FGR:FGR and MICHIGAN:HRC:HRC). Imputed dosage is the expected minor allele count for each imputed genotype based on the posterior imputation probabilities for the genotypes with zero minor alleles (AA), 1 minor allele (Aa) and 2 minor alleles (aa). For the set of heterozygote genotypes investigated here, the correct imputed dosage should be equal to 1. Imputed dosages such as 0.5 that are distant from the set {0,1,2} reflect a high level of uncertainty, and hence inaccurate imputation. No dosage values were observed outside of the range [0-1.2].**

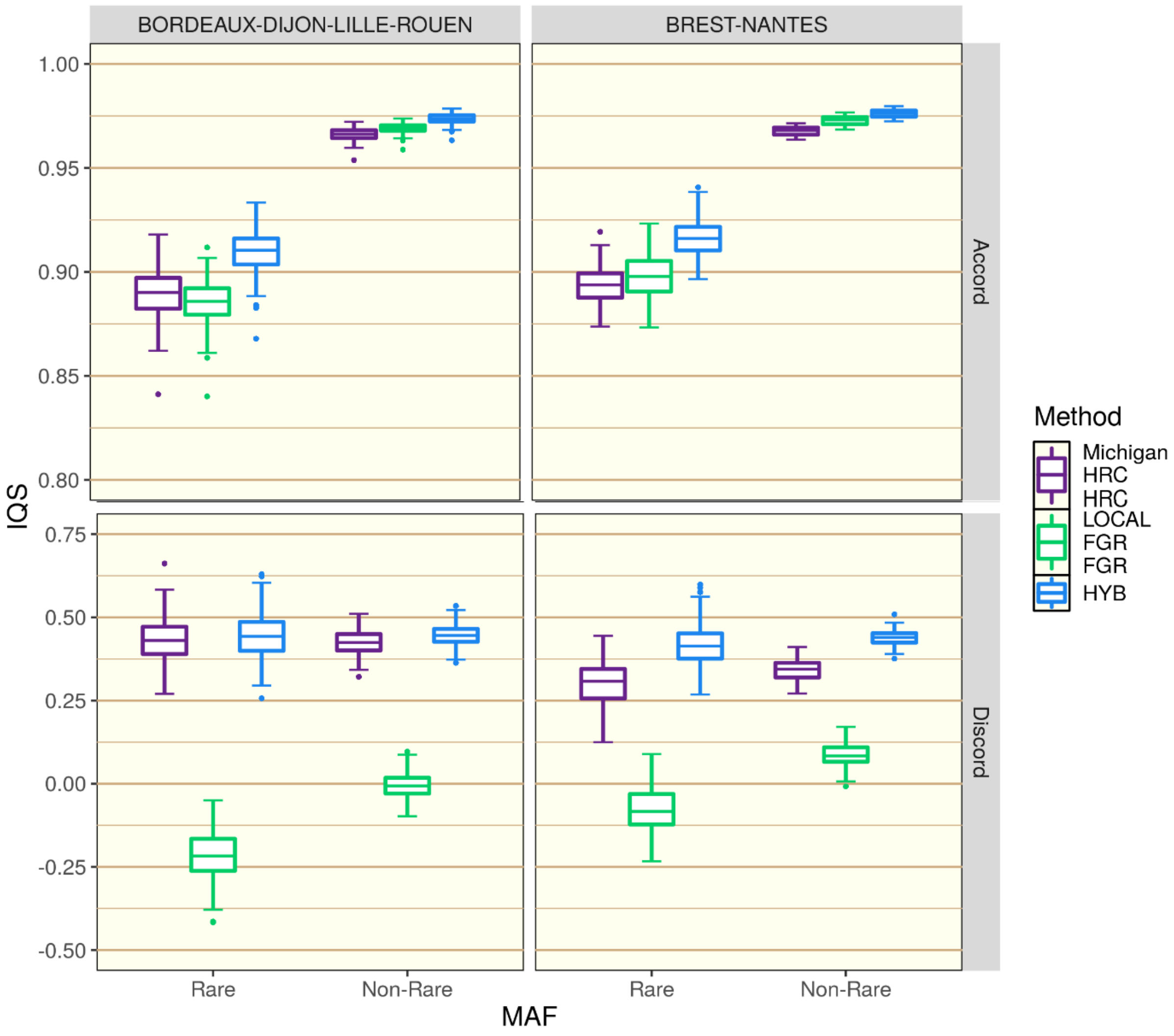


*Supplementary Figure 7. Individual IQS scores calculated for different variant sets. Firstly dichotomising between rare and non-rare variants (MAF below of above 0.01 measured in FrEx) and secondly dichotomising between whether imputation runs MICHIGAN:HRC:HRC and LOCAL:FGR:FGR were in agreement or not (Accord or Discord, respectively) regarding the most likely genotype. Note the difference y-axis ranges on top two and bottom two panels.*

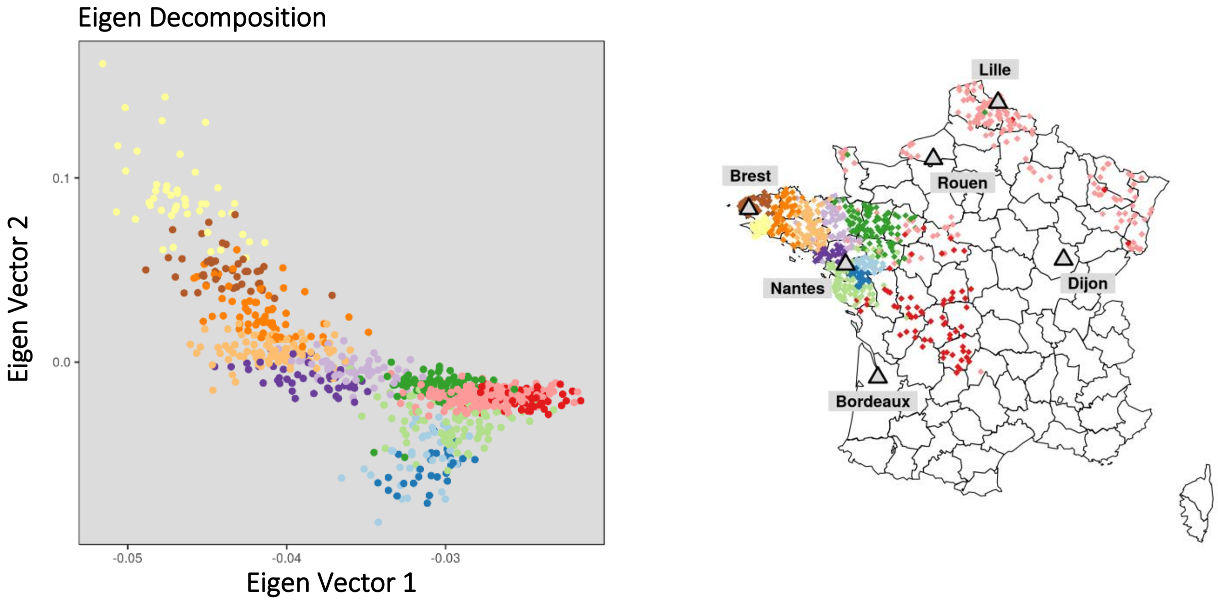


**Supplementary Figure 8. Eigen decomposition analysis of the IBD-sharing matrix in FGR (left). Colours represent the 12 clusters identified by finestructure[6] and described in the main text. The correspondence with geographical locations is also given for comparison (right) where individuals from FranceGenRef are plotted as diamonds.**
